# The virome in adult monozygotic twins with concordant or discordant gut microbiomes

**DOI:** 10.1101/509273

**Authors:** J. Leonardo Moreno-Gallego, Shao-Pei Chou, Sara C. Di Rienzi, Julia K. Goodrich, Timothy Spector, Jordana T. Bell, Youngblut, Ian Hewson, Alejandro Reyes, Ruth E. Ley

## Abstract

The virome is one of the most variable components of the human gut microbiome. Within twin-pairs, viromes have been shown to be similar for infants but not for adults, indicating that as twins age and their environments and microbiomes diverge, so do their viromes. The degree to which the microbiome drives the virome’s vast diversity is unclear. Here, we examined the relationship between microbiome diversity and virome diversity in 21 adult monozygotic twin pairs selected for high or low microbiome concordance. Viromes derived from virus-like particles were unique to each subject, dominated by Caudovirales and Microviridae, and exhibited a small core that included crAssphage. Microbiome-discordant twins had more dissimilar viromes compared to microbiome-concordant twins, and the richer the microbiomes, the richer the viromes. These patterns were driven by the bacteriophages, not eukaryotic viruses. These observations support a strong role of the microbiome in patterning the virome.

## INTRODUCTION

The bulk of the human gut microbiome is composed of a vast diversity of bacterial cells, along with a minority of archaeal and eukaryotic cells. The cellular fraction of the microbiome forms a high density microbial ecosystem (10^11^ −10^12^ per gram of feces (Sender et al., 2016). All of these cells are accompanied by a virome estimated to be in about equal proportion (ranging between 10^9^ to 10^12^ per gram of feces (Castro-Mejía et al., 2015; Hoyles et al., 2014; Ogilvie and Jones, 2017; Reyes et al., 2010). The viral fraction of the human gut microbiome is primarily composed of bacteriophages and prophages, and it also includes rarer eukaryotic viruses and endogenous retroviruses (Breitbart et al., 2003; Minot et al., 2011; Reyes et al., 2010). Currently, the majority of phages have no matches in databases and their hosts remain to be elucidated. Matching phages to their hosts is challenging: for instance, the host of the most common human gut phage, crAssphage, has only recently been identified as *Bacteroides spp*. (Shkoporov et al., 2018; Yutin et al., 2018). In addition to the identification of hosts, other questions remain as to the factors most important in shaping the virome, and how predictive the cellular fraction of the microbiome can be of the virome.

The temporal population dynamics of phages and their hosts might be expected to be linked. Indeed, population oscillations of viruses and their bacterial hosts are described for aquatic systems, where they indicate that viruses play a key role in regulating bacterial populations (Suttle, 2007; Thingstad, 2000; Thingstad et al., 2014; Weitz and Dushoff, 2008). But such patterns of predator/prey dynamics are not typical for the human gut virome and microbiome (for clarity, from here on we use ‘microbiome’ to refer to cellular fraction of the microbiome, e.g., mostly bacterial cells) (Minot et al., 2011; Reyes et al., 2013; Rodriguez-Brito et al., 2010; Rodriguez-Valera et al., 2009). Nonetheless, the virome and microbiome do display some common patterns of diversity across hosts, such as high levels of interpersonal differences and relative stability over time (Reyes et al., 2010). The microbiome tends to be more similar for related individuals compared to unrelated individuals, possibly due to shared dietary habits, which drive similarity between microbiomes (Cotillard et al., 2013; David et al., 2014). In accord, diet has been associated with virome diversity, quite possibly through diet effects on the microbiome (Minot et al., 2011). In infants, twin comparisons have revealed viromes to be more similar between co-twins than between unrelated individuals (Lim et al., 2015; Reyes et al., 2015). This pattern was not observed in adult twins (Reyes et al., 2010) possibly due to divergence of their microbiomes (Reyes et al., 2010). The degree to which the microbiome itself drives patterns of virome diversity across hosts has been difficult to assess due to confounding factors such as host relatedness.

Here, we focus on adult monozygotic (MZ) twin microbiomes to explore further the relationship between microbiome and virome diversity. By studying the viromes of MZ twin pairs, we control for host genetic relatedness. Although MZ twin pairs generally have more similar microbiomes compared to dizygotic (DZ) twin pairs or unrelated individuals, MZ twins nevertheless can display a large range of within-twin-pair microbiome diversity (Goodrich et al., 2014). We previously generated fecal microbiome data for twin pairs from the TwinsUK cohort (Goodrich et al., 2014), and based on this information we selected twin pairs either highly concordant or highly discordant for their microbiomes. We generated viromes from virus-like particles (VLPs) obtained from the same samples from which the microbiomes were derived. Results indicate that microbiome diversity and virome diversity measures are positively associated.

## RESULTS

### Selection of microbiome-concordant and discordant monozygotic twin pairs

We selected twin pairs with a similar body mass index (BMI), whose microbiomes were either concordant or discordant for microbiome between-sample diversity (β-diversity) based on previously obtained 16S rRNA gene data. The adult co-twins in this study did not share a household and we assume that other environmental variability was similar across twin pairs. We determined the degree of concordance or discordance between co-twins’ microbiomes based on three β-diversity distance metrics: Bray-Curtis, weighted UniFrac and unweighted UniFrac **(See Methods)**. As expected, the β-diversity measures were correlated (Pearson pairwise correlation coefficient > 0.4). Based on the distribution of pairwise distance measures, we selected 21 MZ twin pairs from the boundaries of all three distributions (Figure 1A), while maintaining a balanced distribution of age and BMI across the set **(Table S1).** Within the 21 selected twin pairs, the microbiomes of microbiome-concordant co-twins were, as expected, more similar to each other than microbiomes of microbiome-discordant co-twins (p = 6.31 × 10^−12^). The microbiomes of the discordant co-twins differed compositionally at all taxonomic levels, particularly at the phylum level, with Firmicutes and Bacteroidetes, the two dominant phyla, contributing the most to the variation between co-twins (Figure 1B and 1C).

**Figure 1.**
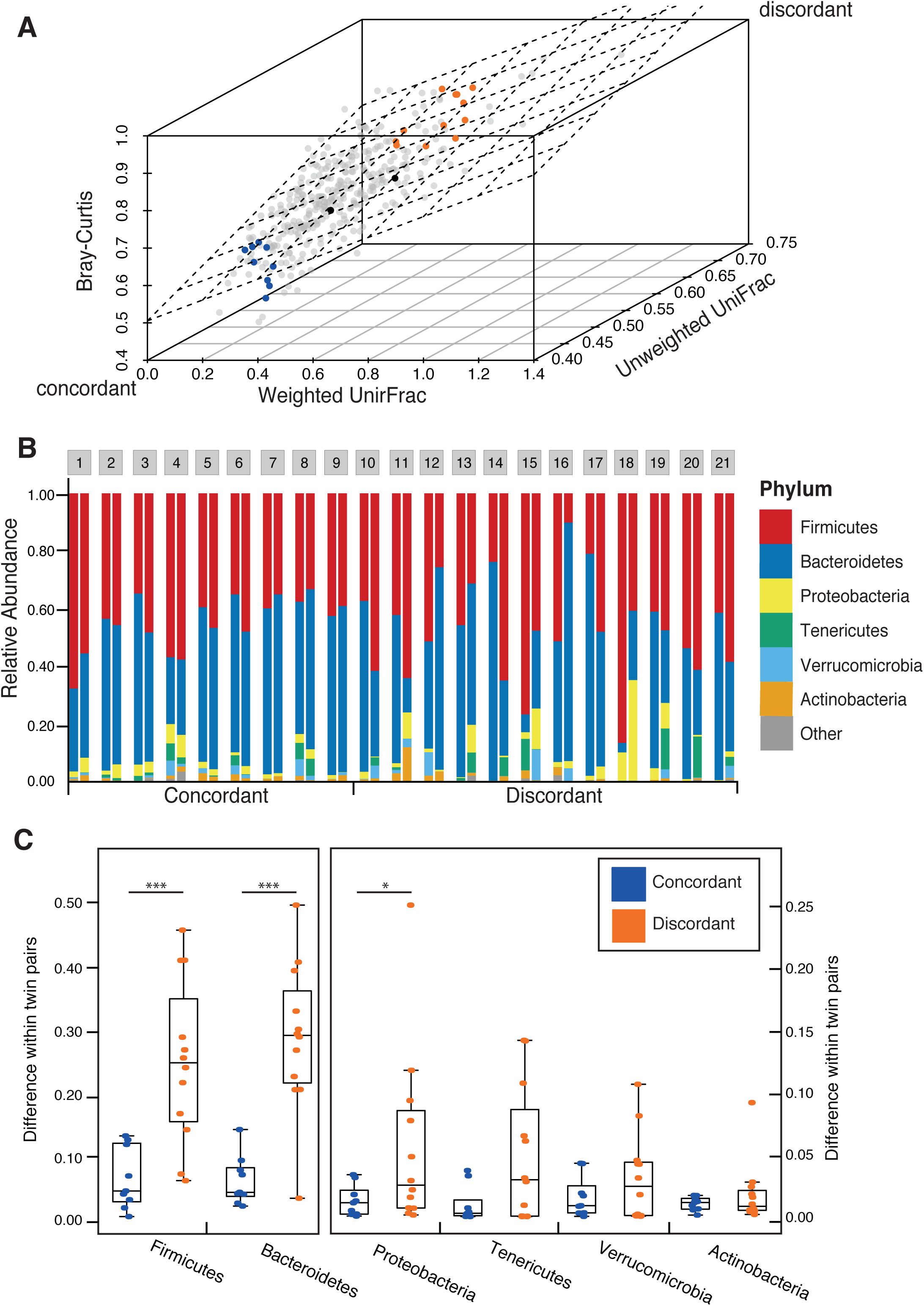
Microbiome discordance in twin pairs. **(A)** The β-diversity measures of the microbiotas of 354 monozygotic twin pairs from a previous study (Goodrich et al., 2014) are shown. Each dot represents the β-diversity of a pair of twins, measured by the weighted UniFrac (x-axis), unweighted UniFrac (z-axis), and Bray-Curtis (y-axis) β-diversity metrics. The three β-diversity metrics are in general correlated (Pearson pairwise correlation coefficient > 0.4). The plane is the least squared fitted plane Bray-Curtis ∼ Weighted UniFrac + Unweighted UniFrac. A subset of twin pairs with concordant microbiotas (blue) and discordant microbiotas (orange) were chosen from the two edges. Black dots indicate the samples used for virome and whole fecal metagenome comparison. **(B)** Comparison of the taxonomic profiles (relative abundance) at the Phylum level for the 21 MZ twin pairs concordant (1-9) or discordant (10-21) for their microbiotas. **(C)** Differences in the relative abundances for the major phyla for concordant (blue points, n=9) and discordant (orange points, n=12) twin pairs. Mann-Whitney’s U test. *** p < 0.0005, * p = 0.055

### Shotgun metagenomes of VLPs

We isolated virus-like particles (VLPs) from the same fecal samples that had been used for 16S rRNA gene diversity profiling **(See Methods)**. DNA extracted from VLPs was used in whole genome amplification followed by shotgun metagenome sequencing **(See Methods)**. A first library (“large-insert-size library”) was selected with an average insert size of 500 bp (34,325,116 paired reads in total; 817,265 ± 249,550 paired reads per sample after quality control) and used for *de novo* assembly of viral contigs. Smaller fragments with an average insert size of 300bp were purified in a second library (“small-insert-size library”) and sequenced. The resulting pair-end reads were merged into 25,324,163 quality filtered longer reads to increase mapping accuracy (602,956 ± 595,444 merged reads per sample) **(See Methods) (Table S2)**.

### Identification of putative bacterial contaminants

Viromes prepared and sequenced from VLPs may be contaminated with bacterial DNA (Roux et al., 2013). However, given that phages are major agents of horizontal gene transfer and that temperate viruses often comprise up to 10% of bacterial genomes in a prophage state, removal of potential bacterial contamination risks also removing viral reads. To assess bacterial DNA contamination, we mapped virome reads against a set of 8,163 fully assembled bacterial genomes. Our strategy consisted of evaluating the coverage along the length of each genome (in bins of 100Kb), and those genomes with a median coverage greater than 100 were considered contaminants. Reads mapping to short regions were considered to be prophages or horizontally transferred genes and retained **(See Methods)** (Figure 2A). Reads mapping to genomes determined to be potential contaminants were removed from further analyses.

**Figure 2.**
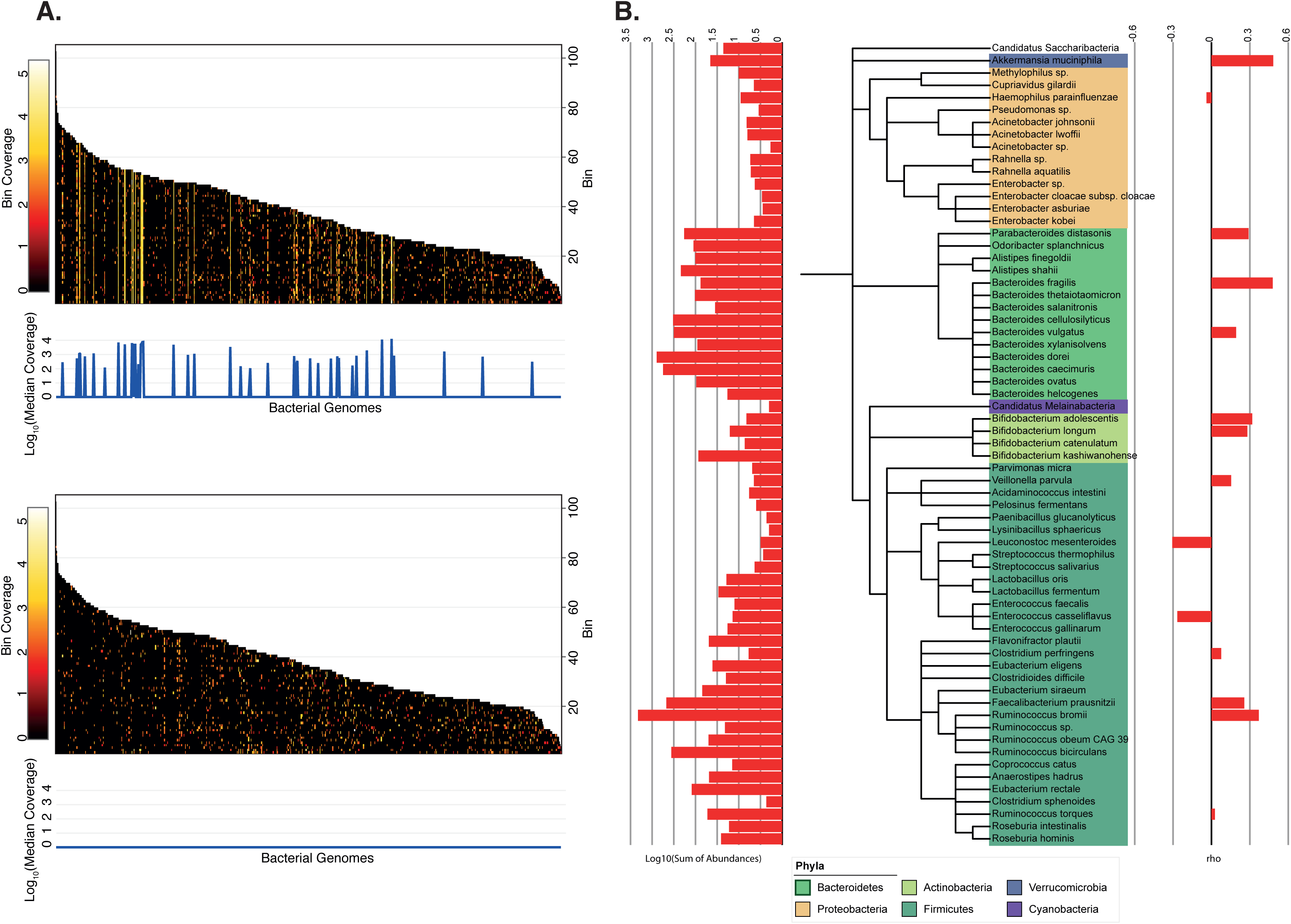
Bacterial contamination in VLP preparations. **(A)** Heatmap of VLP reads from sample 4A mapping to bacterial genomes before and after the removal of reads determined as contaminants. Genomes are sorted by length and split in bins of 100,000 bp. Bacterial genomes with a median coverage greater than 100 were considered as contaminants. **(B)** Cladogram based on the NCBI taxonomy of the 65 genomes identified as contaminants across all VLP extractions. **(Right)** Spearman rank correlation coefficient (rho) between the abundance of the bacterial genomes in the VLP extractions and 16S rRNA gene profile from the microbiome. **(Left)** Total abundance of each bacterial genome added across all individuals.

We identified 65 bacterial genomes as contributing to potential contaminant, with 1.006 ± 1.125% (average ± std) reads per sample mapping to those bacterial genomes **(Table S2)**. The majority (37/68) belonged to the Firmicutes phylum; at the species level, *Bacteroides dorei, B*. *vulgatus, Ruminococcus bromii, Faecalibacterium prausnitzii, B*. *xylanisolvens, Odoribacter splanchnicus and B*. *caecimuris* (in that order) were detectable in at least 50% of the samples **(Table S2)**. If the most abundant bacterial species in the microbiome are the most likely sources of contamination, then the taxonomic composition of the bacterial contaminants should correlate with their corresponding bacterial abundances in the microbiome. However, we observed no significant correlation between the relative abundances of taxa represented in the contaminant DNA and in the microbiomes (Figure 2B).

### Functional profiles support viral enrichment in VLP purifications

To assess the functional content of the viromes, we annotated the “short-insert-size library” raw reads using the KEGG annotation of the Integrated Gene Catalog (IGC) (Li et al., 2014) **(See Methods)**. In line with previous reports (Breitbart et al., 2008; Minot et al., 2011; Reyes et al., 2010), the majority of reads (85.43 ± 5.74%) from our VLP metagenomes mapped to genes with unknown function (Figure 3A).

**Figure 3.**
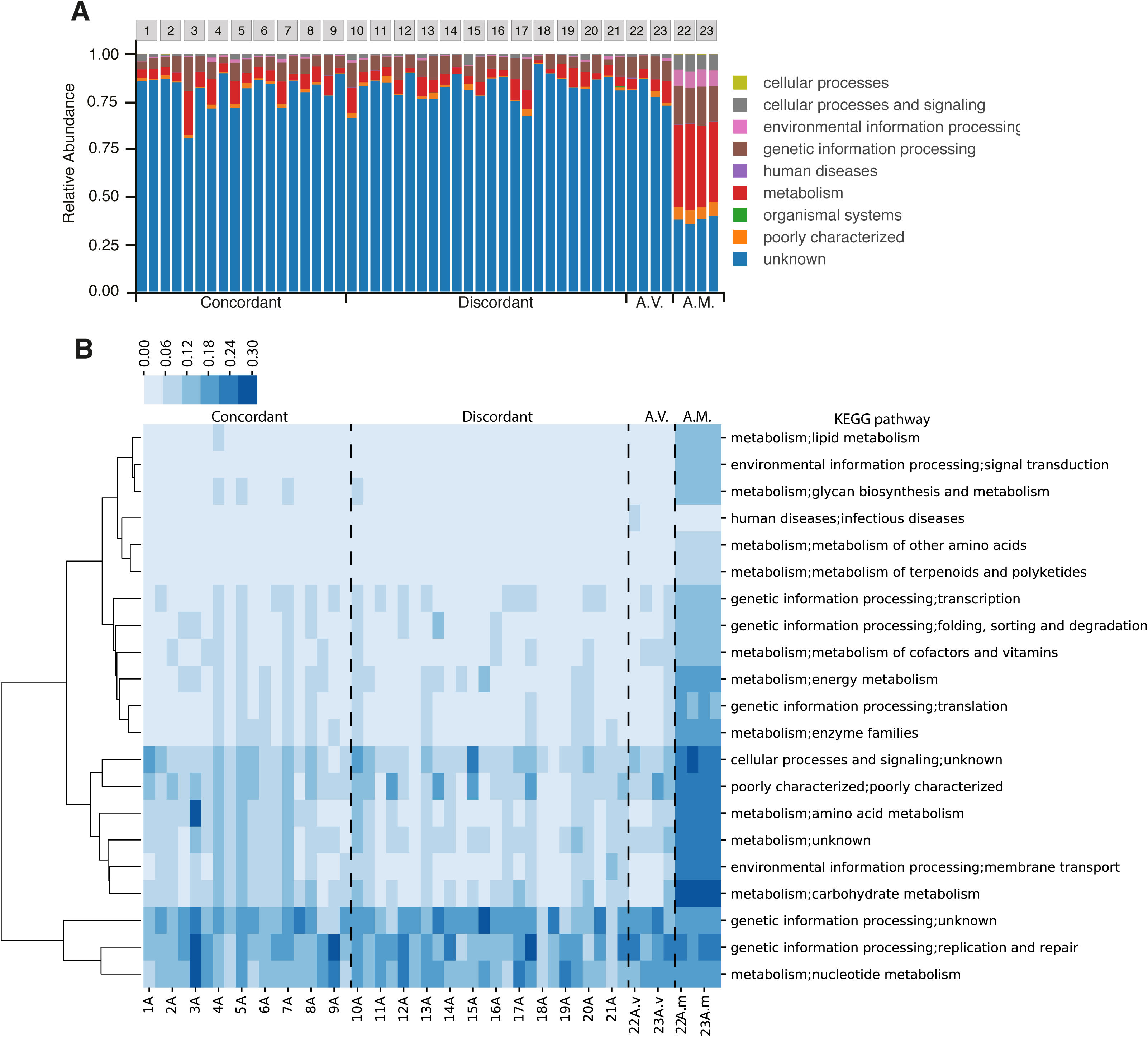
Comparison of the gene content of whole fecal metagenomes and viromes. Relative abundance of KEGG categories in whole fecal metagenomes and viromes. **(A)** The relative abundance of KEGG categories in whole fecal metagenomes and viromes, including all hits to IGC genes, regardless of the annotation. **(B)** Heatmap of the relative abundance of the second level of KEGG categories in whole fecal metagenomes and viromes, excluding the IGC genes with unknown annotation. A.V.: Additional viromes; A.M.: Additional microbiomes (whole genome extractions). Intra-class coefficient (ICC) for A.M. = 0.99; ICC for A.V. = 0.85; ICC concordant-microbiome co-twins = 0.69; ICC discordant-microbiome co-twins = 0.68.

To further verify that sequences were derived from VLPs and not microbiomes generally, we conducted an internal check in which we generated and compared additional metagenomes from VLPs and bulk fecal DNA for an additional 4 individuals (2 twin pairs; Figure 1A). As expected, the functional profiles of viromes and microbiome-metagenomes derived from the same samples were dissimilar. Virome reads that mapped to annotated genes were enriched in two categories: Genetic Information Process (48.87 ± 12.12%) and Nucleotide Metabolism (17.59 ± 8.81%), compared to 24.31 ± 1.28% and 5.47 ± 0.4% for the microbiome-metagenome, respectively (Figure 3B). Most of the other functional categories present in the bacterial metagenomes were essentially absent from the viromes. Furthermore, the functional annotations of the viromes show greater between-sample variability than the microbiomes and a lower intraclass correlation coefficient (Figure 3B).

### Viromes are unique to individuals

We assembled reads from the “large-insert-size library” resulting in a total of 107,307 contigs ≥ 500 nt (max: 79,863 nt; mean 1,186nt ± 1,741; Figure S1). To assess the structure and composition of the viromes, a matrix of the recruitment of reads against dereplicated contigs were built **(See Methods)**. The recruitment matrix included 14,584 contigs that were both long (> 1,300 nt) and well covered (> 5X); these are referred to as ‘virotypes’ (Figure S1). Analysis of the recruitment matrix showed that each individual harbored a unique set of virotypes: 3,415 virotypes (23.41% of total) were present in only one individual; 413 virotypes (2.83%) were present in at least 50% of the individuals; only 18 virotypes (0.1%) were present in all individuals.

### Twins with concordant microbiomes share virotypes

We checked for virotypes shared between twins and observed that co-twins did not share more virotypes than unrelated individuals (p = 0.074). We then assessed microbiome-concordant and discordant twin pairs separately: twins with a discordant microbiome did not share more virotypes that unrelated individuals (p = 0.254), and twins with a concordant microbiome did share more virotypes than unrelated individuals (p = 0.048). Furthermore, we also found that twins with a concordant microbiome shared more virotypes than twins with a discordant microbiome (p = 0.015; Figure S2).

### Bacteriophage dominance of the gut virome

In order to characterize the taxonomic composition of the virome, we attempt to annotated all 66,446 dereplicated and well covered contigs (Figure S1) using a voting system approach that exploited the information in both the assembled contigs and their encoding proteins **(See Methods)**. In addition, we performed a custom annotation on two highly abundant gut-associated bacteriophage families: (i) the crAssphage (Dutilh et al. 2014; Yuting et al. 2018) and (ii) the *Microviridae* families (Székely and Breitbart 2016). For this, we used profile Hidden Markov Models (HMMs) to search for crAssphage (dsDNA viruses) and *Microviridae* (ssDNA viruses) contigs **(See Methods)**.

Using HMMs allowed us to identify distant homologs, which we then incorporated into a phylogenetic tree with known reference sequences to confirm the annotation and better resolve the taxonomy. We annotated 108 contigs (19 crAssphage, 90 *Microviridae*), validated the family assignment of 68 contigs, and assigned a subfamily to 97 contigs without previous subfamily assignment. For the *Microviridae*, only 11 contigs had a previous taxonomic assignment, all belonging to the *Gokushovirinae*: we confirmed these and 23 more as *Gokushovirinae*, 54 as Alpavirinae and 1 contig as *Pichovirinae* (Figure S3A). For the crAssphage, 11 contigs were clustered with the original crAssphage, 3 contigs grouped with the reference Chlamydia phage, and 5 contig grouped with the reference IAS virus (Figure S3B).

After collating the voting system annotation and the HMM annotation, a total of 12,751 contigs (29,62%) were taxonomically assigned (Figure S1). Viromes were dominated by bacteriophages with only 6.42% of contigs annotated as Eukaryotic viruses. As expected, most of the contigs (96.98%) were dsDNA viruses, while only 2.43% of contigs were annotated as ssDNA viruses. Caudovirales was the most abundant Order, with its three main families represented: Myoviridae (20.22 ± 4.83%), *Podoviridae* (10.54 ± 3.27%), and *Siphoviridae* (35.25 ± 7.19%). The crAssphage family constituted on average 13.26% (± 12.24%) of the contigs, reaching a maximum contribution of 55.80% in one virome, and *Microviridae* represented 3.87 ± 2.57% of the viromes. Interestingly, we observed that *Phycodnaviridae* exceeded 1% of average abundance (1.77 ± 1.12%; Figure 4A) and that contigs related to any nucleocytoplasmic large DNA viruses (NCLDV) had a mean relative contribution of 3.99 ± 2.22%. The 18 contigs present in all samples included 10 annotated as crAssphage, 2 annotated as “unclassified Myoviridae”, 2 “unclassified Caudovirales”, 1 classified as *Microviridae*, and 3 unclassified. Within a defined taxonomic profile for each sample, we looked for differences in composition between viromes at all taxonomic levels for concordant and discordant twin-pairs. There were no significant differences between groups for any taxa at the Order and Family levels, including crAssphage and *Microviridae* families (Figure 4B).

**Figure 4.**
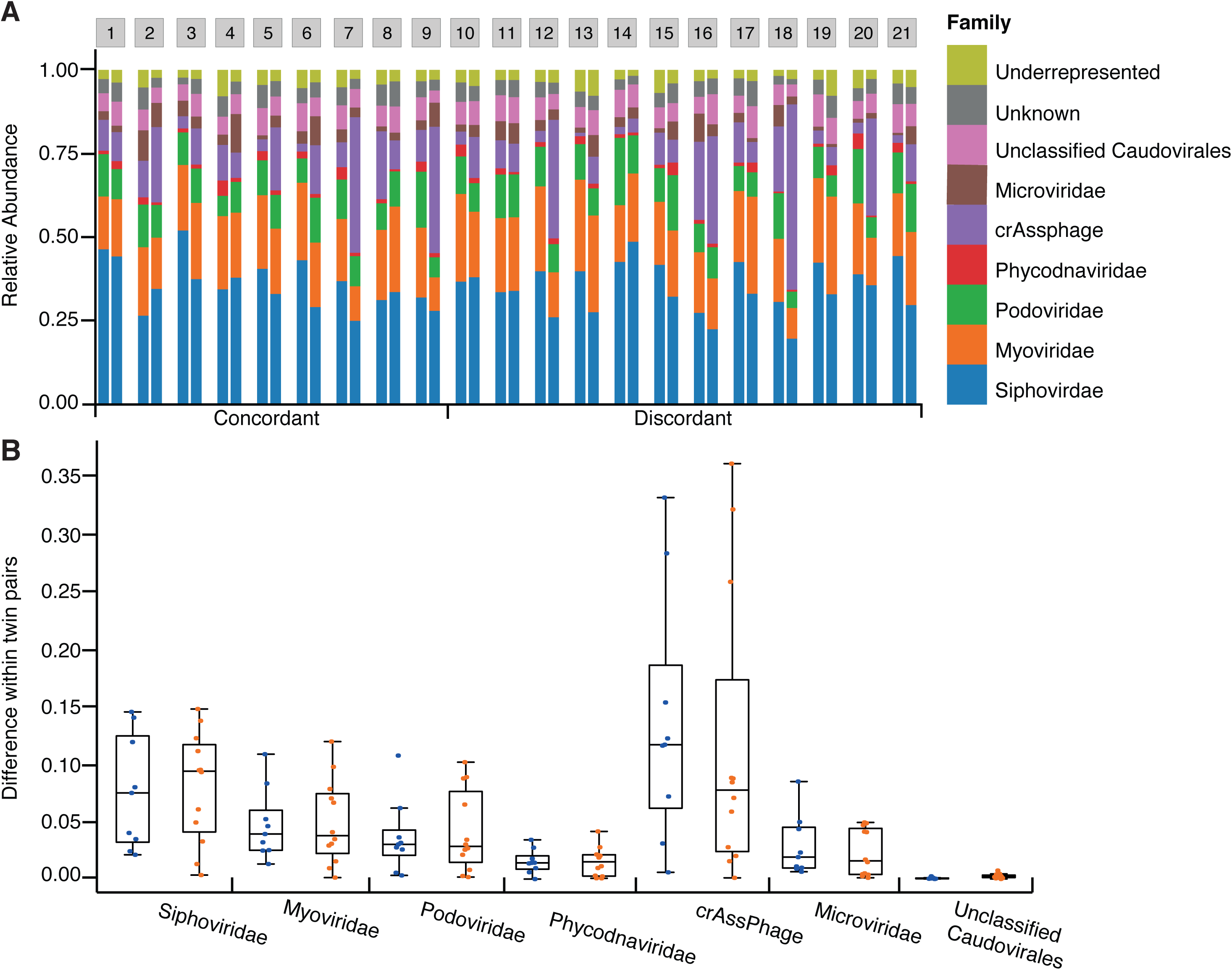
Virome composition. Comparison of the taxonomic profiles at the Family level for the 21 MZ twin pairs concordant (1-9) or discordant (10-21) for their viromes. **(A)** The viral family composition of the MZ twins. **(B)** Differences of the relative abundances of each family for concordant (blue points, n=9) and discordant (orange points, n=12) twin pairs.

We used CRISPR spacer mapping and the microbe-versus-phage (MVP) database (Gao et al., 2018) to predict hosts for virotypes and taxonomically characterized contigs **(See Methods)**. As host annotation was directed to bacteriophages, we did not gain any information for contigs annotated as Eukaryotic viruses. These approaches allowed us to identify putative hosts for 910 contigs. Within these 910 contigs, only one was previously annotated as crAssphage, and ss expected, its host was inferred to be a member of *Bacteroidetes*. In total we identified 1,280 bacterial putative host strains, including 187 species from 87 genera over several phyla; most of them from Firmicutes (92), followed by Bacteroidetes (41) and Proteobacteria (38). The median number of host for each contig was 1 (IQR=1-2) while the median number of phages per host, at the strain level, was 2 (IQR=1-3) (Figure S4).

### Virome diversity correlates with microbiome diversity

To assess the relationship between virome and microbiome diversity, we examined the within-samples diversity (α-diversity) and β-diversity of the viromes using three different layers of information that we recovered from the sequence data: i) virotypes, iii) taxonomically annotated contigs, and iii) annotated genes from short reads (Figure S1).

#### Alpha-diversity

α-diversities of the microbiome and the virome were positively correlated in two of the three layers of information used to test the correlation (virotypes and taxonomy annotated contigs but not genes; Figure 5A). We used annotated contigs to ask about the α-diversity within subgroups of viruses: (ssDNA eukaryotic, dsDNA eukaryotic, ssDNA bacteria and dsDNA bacteria). Our results show that the diversity of eukaryotic viruses does not correlate with the microbiome α-diversity. In contrast, bacteriophages and microbiome α-diversity were positively correlated, for both ssDNA or dsDNA bacterial viruses (Figure 5B).

**Figure 5.**
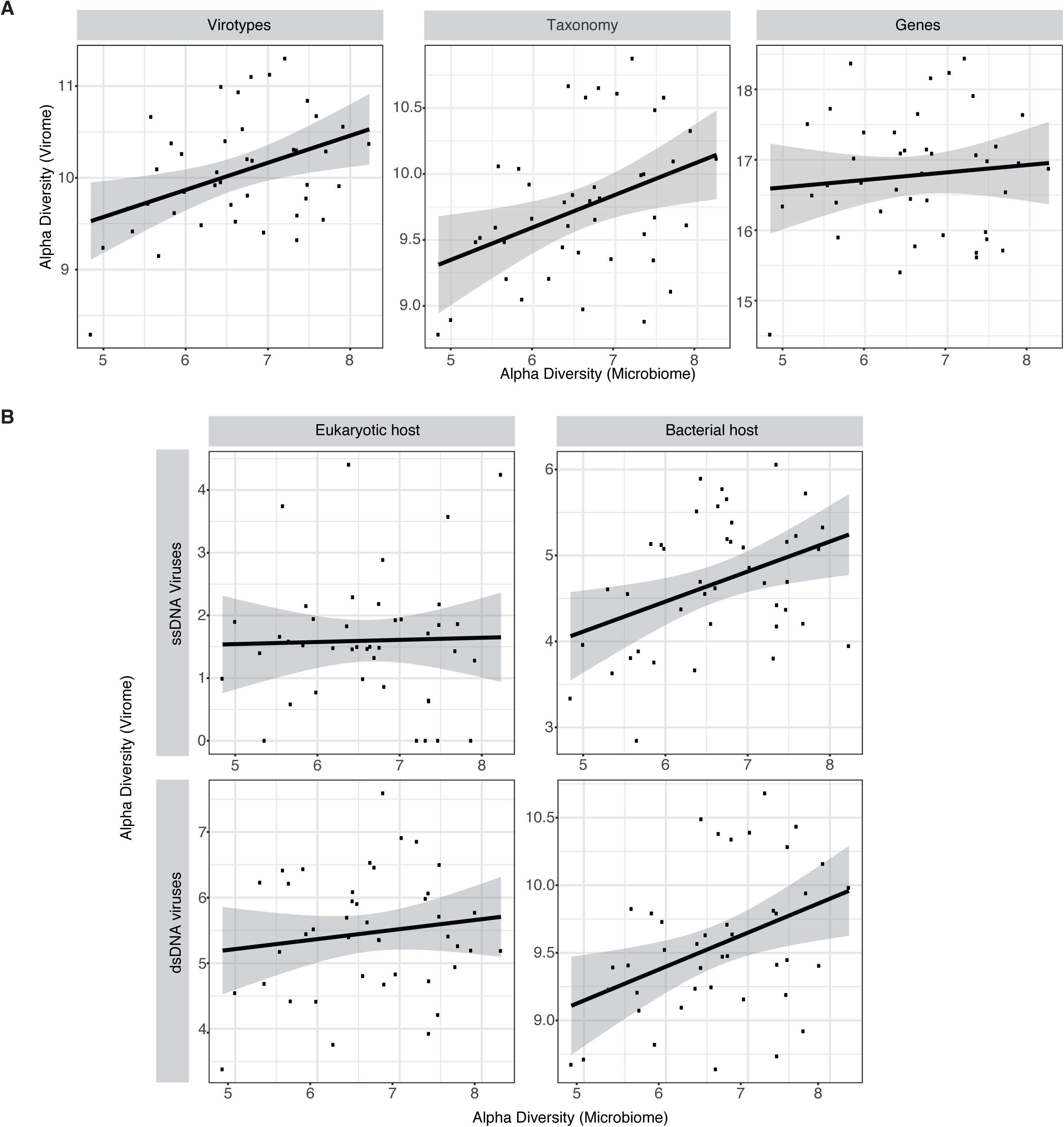
Bacteriophages diversity correlates with microbiome diversity but eukaryotic viruses diversity do not. **(A)** Correlation of Shannon α-diversity of viromes to Shannon α-diversity of microbiomes (n=42). **i) Virotypes:** Pearson correlation coefficient = 0.406, m = 0.3, p = 0.007, R^2^ = 0.165; **ii) Taxonomy:** Pearson correlation coefficient = 0.389, m = 0.25, p = 0.010, R^2^ = 0.151; iii) **Genes:** Pearson correlation coefficient = 0.105, m = 0.11, p = 0.506, R^2^ = 0.011 **(B)** Correlation of the Shannon α-diversity of the virome, calculated from contigs annotated as ssDNA eukaryotic viruses, ssDNA phages, dsDNA eukaryotic viruses, and dsDNA phages, to Shannon α-diversity of the microbiome (n=42). **ssDNA eukaryotic viruses:** Pearson correlation coefficient = 0.027, m = 0.034, p = 0.863, R^2^ = 0.000751; **ssDNA bacteriophages:** Pearson correlation coefficient = 0.394, m = 0.35, p = 0.009, R^2^ = 0.155; **dsDNA eukaryotic viruses:** Pearson correlation coefficient = 0.143, m = 0.15, p = 0.368, R^2^ = 0.020; **dsDNA bacteriophages:** Pearson correlation coefficient = 0.400, m = 0.25, p = 0.008, R^2^ = 0.16.

#### Beta-diversity

We observed that concordant twins had lower virome β-diversity compared to discordant twins using Hellinger distances (Figure 6); the mean binary Jaccard distance and Bray-Curtis dissimilarity of viromes also showed the same trend (Figure S5A and S5B). Similar to what we observed with α-diversity, regardless of the layer of information used, the mean Hellinger distance of viromes within MZ twin pairs with concordant microbiomes was significantly lower than that of MZ twin pairs with discordant microbiomes (p < 0.04, Mann-Whitney’s U test) (Figure 6). Furthermore, a similar significant positive correlation was observed between microbiome and virome β-diversity when using the annotated contigs. This relationship was driven by the bacteriophages (p = 0.009, Mann-Whitney’s U test), but not the eukaryotic viruses (p = 0.243, Mann-Whitney’s U test).

**Figure 6.**
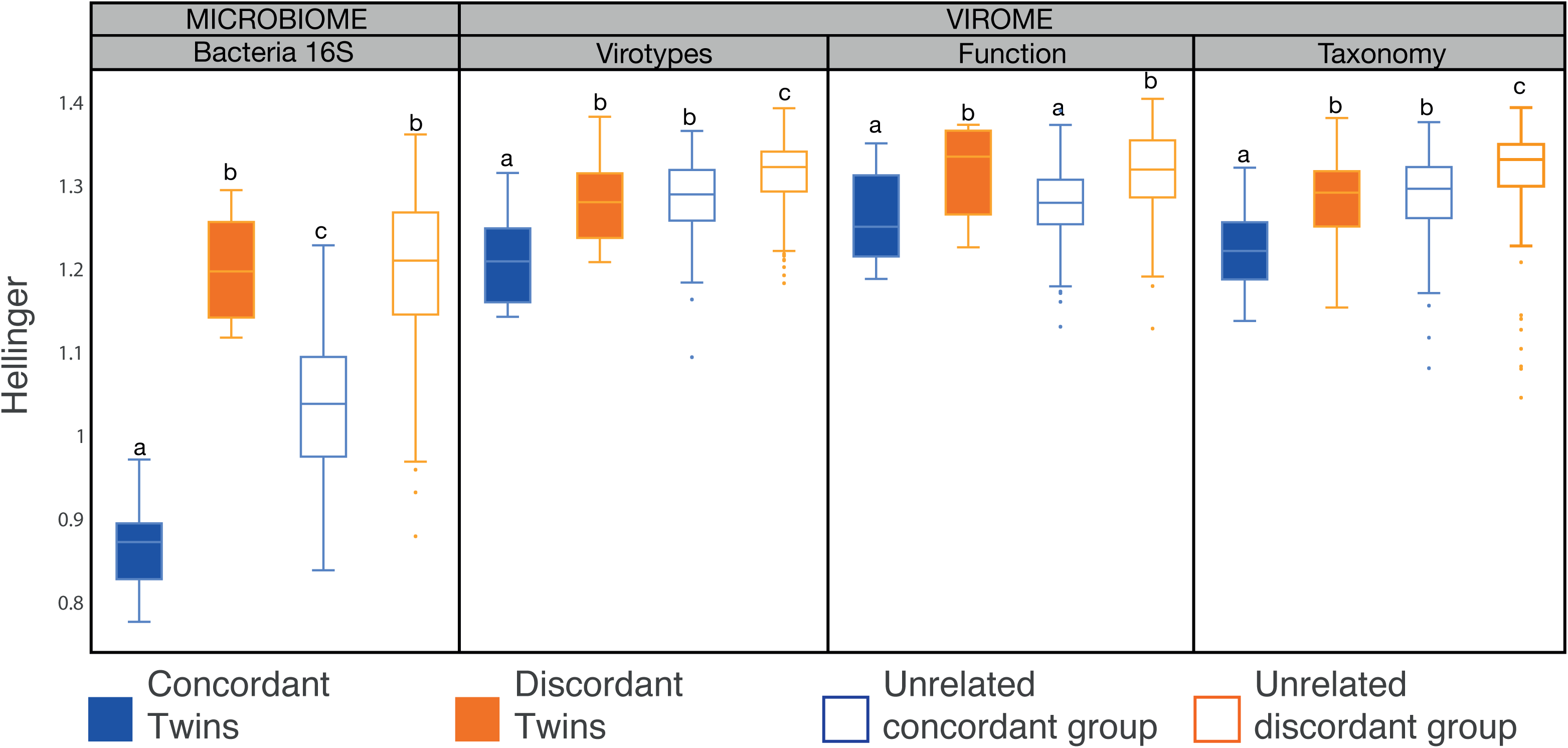
Virome Beta-diversity patterns mirror microbiome Beta-diversity. Box plots show the distribution of Hellinger distances for microbiomes and viromes, according to the three different layers of information recovered (virotypes, function, and taxonomy), for concordant co-twins (blue, n=9), discordant co-twins (orange, n=12), unrelated samples within the concordant co-twins (blue edges, n=144), and unrelated samples within the discordant co-twins (orange edges, n=264). Significant differences between means (Mann-Whitney’s U test, p < 0.020) are denoted with different letters.

Finally, we compared the virome and microbiome pairwise distances among related (co-twins) and unrelated individuals. The pairwise distance matrices showed a positive correlation between virome and microbiome β-diversity measures not only within twin pairs (Pearson correlation coefficient > 0.50) but also generally across all individuals (Pearson correlation coefficient > 0.25; p < 0.003, Mantel test; Figure S5C). These results show that regardless of genetic relatedness between hosts, individuals with more similar microbiomes harbour more similar viromes.

## DISCUSSION

Co-twins, like other siblings, generally have more similar gut microbiomes within their twinships compared to unrelated individuals (Lee et al., 2011; Palmer et al., 2007; Tims et al., 2013; Turnbaugh et al., 2009; Yatsunenko et al., 2012). Moreover, MZ twins have overall more similar microbiomes than DZ twins, although at a whole-microbiome level this effect is small and primarily driven by a small set of heritable microbiota (Goodrich et al., 2014, 2016). Within a population of MZ twin pairs, however, the range of within-twin pair differences in the microbiomes can be as great as for DZ twins (Goodrich et al., 2014). We took advantage of the large spread in β-diversity for MZ co-twins to select co-twins that were either highly concordant or discordant for their gut microbiomes. Our analysis of their viromes showed that despite the high variation in the gut viromes between individuals, and regardless of host relatedness, the more dissimilar their microbiomes, the more dissimilar their viromes. This pattern was driven by the bacteriophage component of the virome.

Here, by choosing MZ twins from a distribution of divergence in the microbiome, we removed host genetic relatedness as a variable. Previous studies of the viromes and microbiomes of infant twin pairs showed that the microbiomes and viromes of co-twins were more similar than those of unrelated individuals, suggested shared host genotype and/or environment were key (Lim et al., 2015; Reyes et al., 2015). In contrast, an early study of the virome of adult twins showed that adult co-twins did not have more similar viromes than unrelated individuals (Reyes et al., 2010); however, in light of the current study’s results, this was likely a power issue. Indeed, in our dataset we observed that regardless of whether twins were concordant or discordant for their microbiomes, co-twins had more similar viromes (virotypes and taxonomy) than unrelated individuals.

The previously reported greater virome similarity in young compared to adult twins has been related to the fact that infants have a greater shared environment compared to adult twins (Lim et al., 2015), particularly in terms of their diet. Minot et al., have also shown that individuals on the same diet have more similar gut viromes than individuals on dissimilar diets (Minot et al., 2011). It is well established that diet is a strong driver of daily microbiome fluctuation (Claesson et al., 2012; David et al., 2014; De Filippo et al., 2010; Wu et al., 2011), so the effect of diet on the virome is likely mediated by the microbiome. However, we did not control for diet, so it is possible that the microbiome discordance that we observe was caused by co-twins eating differently around the time of sampling. Regardless of what underlies the variance in microbiome concordance, it is strongly associated with virome concordance.

The relationship between virome richness and microbiome richness had not previously been directly addressed in adults. We observed that the α-diversity of the microbiome and the virome were positively correlated using two of the three layers of information describing virome diversity. Specifically, this pattern was observed for virotypes and taxonomy but not for genes. However, since virome genes were observed to be enriched in only two categories, Genetic Information Processing and Nucleotide Metabolism, we would not expect differences in diversity of virome genes between subjects. The taxonomic annotation layer showed that the bacteriophage component of the virome, not the eukaryotic viruses, was driving this α-diversity correlation pattern.

The positive relationship between virome and microbiome α-diversity suggests that a greater availability of hosts drives a greater availability of viruses. These observations are in accordance with “(Minot et al., 2013; Reyes et al., 2010), which posits that in a (Minot et al., 2013; Reyes et al., 2010) (Knowles et al., 2016). Indeed, longitudinal studies of the human gut virome have reported genes associated with lysogeny, low mutation rate over time in temperate-like contigs, and long-term stability of the virome, suggesting preference for a lysogenic cycle (Minot et al., 2013; Reyes et al., 2010). Nevertheless, phage predation has been acknowledged as an important factor for the maintenance of highly diverse and efficient ecosystems (Rodriguez-Valera et al., 2009) and may play a role in the maintenance of diversity in a rapidly changing ecosystem as the human gut (David et al., 2014). Short scale time-series analyses of virome-microbiome interactions, along with a better understanding of the lysogenic-lytic switch in viral reproduction, would help to interpret the observed patterns in the human gut virome.

The composition of the viromes described here was similar to what has been previously reported for adult fecal viromes (Minot et al., 2011, 2013; Reyes et al., 2010) but stands in contrast to what has been observed in babies (Lim et al., 2015). From the annotated fraction of the virome, the order *Caudovirales* and its families *Siphoviridae, Myoviridae*, and *Podoviridae*, along with crAssphage, were the dominant phages in all samples. Manrique *et al*. have summarized the phage colonization of the infant gut as follows: the eukaryotic viruses first dominate the newborn gut, followed by the *Caudovirales*, and by 2.5 years of age the *Microviridae* start to dominate (Manrique et al., 2017). We did observe abundant *Microviridae* in our sample set, but the Caudovirales were the dominant group. Age was not related to patterns of diversity in the set of adult subjects studied here.

Despite the high diversity and uniqueness of each virome described here, we nonetheless recovered a core virome among the subjects: 18 contigs were present in all samples. More than half of these contigs were annotated as crAssphage, consistent with recent reports that this phage is widespread (Dutilh et al., 2014; Manrique et al., 2016; Yarygin et al., 2017). Other shared virotypes in our dataset were classified as *Myoviridae* and *Microviridae*. We also recovered contigs mapping to representative families of the nucleocytoplasmic large DNA viruses (NCLDV), *Phycodnaviridae* and *Mimiviridae*. These types of viruses are increasingly reported as members of the human gut virome (Colson et al., 2013; Halary et al., 2016). A core set of bacteriophages consisting of nine representatives, including crAssphage, has previously been reported for the human gut (Manrique et al., 2016). Widely shared virotypes may indicate the wide sharing of specific hosts between individuals, or that these viruses have a broad host range within the human microbiome.

Our use of the HMMs to annotate viral contigs allowed a deep exploration into the taxonomic content of the virome. We annotated a diversity of contigs beyond what was revealed from comparisons to public databases, and also confirmed those annotations. Because each type of virus (*e*.*g*., family) requires its own HMM, we applied this method to a few key groups. When applied to the crAssphage, the HMM retrieved contigs that grouped only with sequences derived from fecal viromes and not with sequences from other environments (e.g., terrestrial or marine). This suggests that although crAssphage is a diverse group of bacteriophages, its diversity in the human gut is restricted to sequences related to the reference crAssphage genome (Dutilh et al., 2014), the IAS virus reference (Shkoporov et al., 2018), or *Chlamydia* bacteriophage (Yutin et al., 2018). We also applied HHM to the family *Microviridae*, which are single strand DNA bacteriophages. We were able to confirm the presence of diverse members of *Gokushovirinae* and Alpavirinae subfamilies.

Although there is evidence that described Alpavirinae genomes constitute a third group of the Microviridae family (Krupovic and Forterre 2011; Roux et al. 2012), they correspond to prophages, which makes it difficult to integrate them into the taxonomy of the International Committee on Taxonomy of Viruses (ICTV), thus, no contigs were annotated as Alpavirinae prior to application of the HMM profiles.

For each taxonomic group of viruses, there is a corresponding set of bacterial hosts. From the 16S rRNA gene diversity data we used to select the twin pairs, it is clear which bacteria phyla contribute the most to the differences in the microbiomes of concordant and discordant twins. But unlike for bacteria, we were not able to discern such clear patterns by order or family in the virome. Indeed, most of the bacteriophage diversity is grouped in just one order, *Caudovirales*, and its three families *Myoviridae, Podoviridae* and *Siphoviridae*. Representatives of these families can infect unrelated hosts (Barylski et al., 2017). As such, we wouldn’t necessarily expect specific orders or families of viruses to show the patterns observed in the bacterial phyla.

Finally, we noted an interesting pattern of complete bacterial genome coverage for select bacteria in the genomes. As these putative contaminants were not the most abundant members of the microbiome, they are unlikely to represent random contamination of bulk DNA. Why certain bacterial genomes showed such high coverage is unclear. One possibility is that we are observing the host species range of transposable phages. Phages such as the Mu phage randomly integrate into the host genome (Taylor, 1963), amplify by successive rounds of replicative transposition, and then can package any section of their host’s genome (Hulo et al., 2011; Toussaint and Rice, 2017). Intriguingly, several of the contaminants detected here (*e*.*g*., *B*. *vulgatus, B*. *dorei, F*. *prausnitzii* and *B*. *thetaiotaomicron*) have also been reported as contaminants in other human gut virome studies (Minot et al., 2011; Roux et al., 2013), which could indicate host-specificity of Mu phages. Alternative explanations include vesicle production, gene transfer agents and/or generalized transduction processes (Biller et al., 2014; McDaniel et al., 2010; Minot et al., 2011). Further comparisons of whole bacterial genomes recovered in diverse virome datasets may help shed light on their source, particularly if the same bacterial species are recovered across multiple studies.

### Prospectus

Our results show that gut microbiome richness and diversity correlate to virome richness and diversity, and vice-versa. The mechanics underlying this association remain to be resolved for the human gut. That the two are coupled may be useful to take into consideration when designing future studies of the virome and factors affecting. Baseline microbiome diversity may be important to balance between groups, for instance, prior to assessing the diversity of the virome.

## METHODS

### Selection of concordant and discordant monozygotic twin pairs

From 16S rRNA gene diversity previously measured for 354 monozygotic twin pairs whose fecal samples were received between January 28th 2013 and July 14th 2014 (Goodrich et al., 2014), we selected 11 concordant and 13 discordant MZ co-twins based on three microbiota β-diversity distances within twin pairs: unweighted UniFrac, weighted UniFrac (Lozupone et al., 2007) and Bray-Curtis (Bray and Curtis, 1957). The twins pairs in the the concordant and the discordant groups were selected to be balanced between those two groups for age, BMI, and BMI difference within a twin pair **(TableS1)**. Twins within the concordant group ranged in age from 23 to 77 years old and included 5 men and 4 women, while those in the discordant group ranged in age from 29 to 81 years old with 5 men and 7 women.

### Isolation of virus-like particles (VLPs) from human fecal samples

VLP isolation procedures were based on the protocol described by (Gudenkauf et al., 2014) and Minot *et al*. *(Minot et al*., *2013*). For VLP isolation, ∼0.5 g of fecal sample was resuspended by vortexing for 5-10 minutes in 15 ml PBS, previously filtered through 0.02 µm filter (Whatman). The homogenates were centrifuged for 30 min at 4,500 *x*g, and the supernatant was filtered through 0.22 µm polyethersulfone (PES) Express Plus Millipore Stericup (150 ml) to remove cell debris and bacterial-sized particles. The filtrate was then concentrated on a Millipore Amicon Ultra-15 Centrifugal Filter Unit 100K to ∼1 ml. The concentrate was transferred to 5 Prime Phase Lock Gel and incubated with 200 µl chloroform for 10 min at room temperature. After being centrifuged for 1 min at 15,000 *x*g, the aqueous layer was transferred to a new microcentrifuge tube, and was treated with Invitrogen TURBO DNase (14 U), Promega RNase One (20 U) and 1 µl Benzonase Nuclease (E1014 Sigma Benzonase^®^ Nuclease) at 37 °C for 3 hr (Gudenkauf and Hewson, 2016; Reyes et al., 2012). After incubation, 0.04 volumes 0.5 M EDTA was added to each sample. The sample was then stored at −80 °C before further processing.

### Viral DNA shotgun sequencing

The viral DNA was extracted with PureLink® Viral RNA/DNA Mini Kit from Invitrogen™. Each viral DNA sample was then amplified using GenomePlex® Complete Whole Genome Amplification (WGA2) Kit from Sigma-Aldrich (Gudenkauf and Hewson, 2016). Two blank controls were included in this step, but very low yield precluded library construction. The amplified product was then fragmented with Covaris S2 Adaptive Focused Acoustic Disruptor with the parameters set as follows: the duty cycle set at 10%, cycle per burst 200, intensity 4 and duration 60 seconds. Each viral sequencing library was prepared following Illumina TruSeq DNA Preparation Protocol with one unique barcode per sample. All barcoded libraries were pooled together. Half of the pool was size selected by BluePippin (Sage Science, Beverly, MA, USA) to enrich fragments with longer inserts (425 bp to 875 bp including the adapters). Both pools, the “large-insert-size library” and the “short-insert-size library”, were sequenced in independent lanes on an Illumina HiSeq 2500 instrument, operating in Rapid Run Mode with 250 bp paired-end chemistry at the Cornell Genomics facility.

### Whole fecal metagenome shotgun sequencing

The genomic DNA was isolated from an aliquot of ∼100 mg from each sample using the PowerSoil® - htp DNA isolation kit (MoBio Laboratories Ltd, Carlsbad, CA). Each sequencing library was then prepared following Illumina TruSeq DNA Preparation Protocol with 500 ng DNA using the gel-free method, 14 cycles of PCR, and with one unique barcode per sample. Sequencing was performed on an Illumina HiSeq 2500 instrument in Rapid Run mode with 2 × 150 bp paired-end chemistry at the Cornell Biotechnology Resource Center Genomics Facility.

### Assessment of Bacterial Contamination

A set of 8,163 finished bacterial genomes was retrieved from the NCBI FTP on 21 February 2017. Reads per sample were mapped against this bacterial genomes dataset using Bowtie2 v.2.2.8 (Langmead and Salzberg, 2012) with the following parameters: --local --maxins 800 - k=3. Genome coverage per base was calculated considering only reads with a mapping quality above 20 using *view* and *depth* Samtools commands v.1.5 (Li et al., 2009). Next, genome coverage was averaged for 100Kbp bins. We observed that evenly covered genomes had a median bin coverage of at least 100; those genomes with a median bin coverage greater than 100 were considered as contaminants. The reads mapping to those genomes were removed. Bacterial genomes can have one or more prophage(s) in their genomes (Munson-McGee et al., 2018) bursting events of those prophages can occur, generating several VLPs. As a conservative measure to avoid the loss of reads originating from prophages and not the bacterial genome *per se*, bins with a coverage over three standard deviations of the bacterial mean coverage were also identified and catalogued as prophages-like regions. Reads mapping to potential contaminant genomes were tagged as “contaminants” and removed from further analysis while reads mapping to high coverage bins were tagged as “possible prophages”.

A matrix of the abundance of each potential contaminant per sample was built using an in-house Python script and normalized by RPKM. In parallel, from Goodrich *et al*. data (Goodrich et al., 2014), the relative abundance of each OTU was recovered and summarized at the species level using summarize_taxa.py qiime script. The Spearman rank order correlation between relative abundances of contaminants and their corresponding 16S rRNAs data was calculated for species in both sets.

### Functional profiles

The joined and trimmed reads from the “short-insert-size library” were mapped onto Integrated Gene Catalogs (IGC), an integrated catalog of reference genes in the human gut microbiome (Li et al., 2014) by BLASTX using DIAMOND v.0.7.5 (Buchfink et al., 2015) with maximum e-value cutoff 0.001, and maximum number of target sequences to report set to 25.

After the mapping onto IGC, an abundance matrix was generated using an in-house Python script. The matrix was then annotated according to the KEGG annotation of each gene provided by IGC. The annotated abundance matrix was rarefied (subsampling without replacement) to 2,000,000 read hits per sample. The KEGG functional profile was then generated using QIIME 1.9 (Quantitative Insights Into Microbial Ecology) (Caporaso et al., 2010) using the command summarize_taxa_through_plots.py. The Intraclass Correlation Coefficient of the functional profiles for each group (additional microbiomes, additional viromes, viromes of concordant-microbiome samples and viromes of discordant-microbiome samples) was calculated using the Psych R package.

### *De-novo* assembly

Reads from the “large-insert-size library” that remain paired (forward and reverse) after the trimming step were assembled using Integrated metagenomic assembly pipeline for short reads (InteMAP) (Lai et al., 2015) with insert size 325 bp ± 100 bp. Each sample was assembled separately. After the first run of assembly, all clean reads were mapped to the assembled contigs using Bowtie2 v.2.2.8 (Langmead and Salzberg, 2012) with the following parameter: --local --maxins 800. The pairs of reads that aligned concordantly at least once were then submitted for the second run of assemble by InteMAP. Contigs larger than 500 bp from all samples were pooled together and compared all vs all, using an in-house Perl script, on the comparison file it was possible to identify potential circular genomes, and dereplicate contigs that were contained in over 90% of their length within another contig.

In order to build an abundance matrix, the recruitment of reads to the dereplicated metagenomic assemblies was used implementing a filter of coverage and length as recommended in Roux *et al*. (Roux et al., 2017). With this in mind, reads (not tagged as contaminants in the previous step) were mapped to dereplicated contigs using Rsubread v.1.28.0 (Liao et al., 2013). Mapping outputs were parsed using an in-house Python script into an abundance matrix that was normalized by reads per kilobase of contig length per million sequenced reads per sample (RPKM) and transformed to *Log*_*10*_*(x+1)*, being *x* the normalized abundance. Contigs with a normalized coverage bellow 5x were excluded. Finally, to virotypes, a filter on contig length was applied. A length threshold was chosen as the elbow of the decay curve generated when plotting the number of contigs as a function of length, which occurred at a length of 1,300 bp.

### HMM annotation

Independent HMM profiles were built to identify crAss-like contigs and Microviridae contigs. To build the HMM-crAsslike profile, sequences for the Major Capsid Protein (MCP) of the proposed crAss-like family (Yutin et al., 2018) were retrieved from ftp.ncbi.nih.gov/pub/yutinn/crassphage_2017/. Multiple sequence alignments (MSA) were done using MUSCLE v.3.8.31(Edgar, 2004) and inspected using UGENE v.1.31.0 (Okonechnikov et al., 2012); positions with more than 30% of gaps were removed. Finally, the HMM-crAsslike profile was built using *hmmbuild* from the HMMER package v.3.1b2 (http://hmmer.org/) (Eddy, 1998). For the Microviridae case, all HMM-profiles for the viral protein 1 (VP1) developed by Alves *et al*. (Alves et al., 2016) were adopted.

Predicted proteins of the assembled contigs were queried for matching the HMM-profiles using *hmmsearch* (Eddy, 1998). Matching proteins with an e-value below 1 × 10^−5^ were considered as true homologs but only proteins between the size rank of the reference proteins (crAsslike MCP: 450-510 residues; Microviridae: 450-800 residues), a coverage of at least 50% and a percentage of identity of at least 40% to at least one reference sequence were used for further analysis. Coverage and identity percentage were determined making a BLASTp of the true homologues against the reference sequences.

True homologues passing the filters mentioned above were used in phylogenetic analysis. Reference and homologous sequences were aligned using MUSCLE v.3.8.31 and sites with at least 30% of gaps were removed using UGENE v.1.31.0. A maximum-likelihood (ML) phylogenetic analysis was done using RAxML v.8.2.4 (Stamatakis, 2014), the best evolutive model was obtained with prottest v.3.4.2 (Darriba et al., 2011) and support for nodes in the ML trees were obtained by bootstrap with 100 pseudoreplicates.

### Taxonomic profiles

To infer the taxonomic affiliation of the assembled VLPs, genes were predicted from all assembled contigs larger than 500 bp using GeneMarkS v.4.32 (Besemer et al., 2001). The amino acid sequence of the predicted genes was then used in a BLASTp search against the NR NCBI viral database using DIAMOND v.0.7.5 (Buchfink et al., 2015) with maximum e-value cutoff 0.001 and maximum number of target sequences to report set to 25. Using the BLASTp results, the taxonomy of each gene was assigned by the lowest-common-ancestor algorithm in MEtaGenome ANalyzer (MEGAN5) v.5.11.3 (Huson et al., 2011) with the following parameters: Min Support: 1, Min Score: 40.0, Max Expected: 0.01, Top Percent: 10.0, Min-Complexity filter: 0.44. Independently, the taxonomy annotation of each contig was obtained using CENTRIFUGE v.1.0.4 (Kim et al., 2016) against the NT NCBI viral genomes database. The final taxonomic annotation of each contig was then assigned using a voting system where the taxonomic annotation of each protein and the CENTRIFUGE annotation of the contig were considered as votes. With all the possible votes for a contig, an N-ary tree was build and the weight of each node was the number of votes including that node. The taxonomic annotation of a contig will be the result of traverse the tree passing through the heaviest nodes with one consideration: if all children nodes of a node have the same weight the traversing must be stopped. The taxonomic profile was considered as a subset of the recruitment matrix containing all contigs annotated either by the voting system or annotated through the HMM profiles (see above).

### Prediction of phage-host interaction

Clustered Regularly Interspaced Short Palindromic Repeats (CRISPRs) were identified using the PilerCR program v.1.06 (Edgar, 2007) from the same set of 8,163 bacterial used to asses the bacterial contamination. Spacers within the expected size of 20 bp and 72 bp (Horvath and Barrangou, 2010) were used as queries against virotypes and taxonomically annotated contigs using BLASTn (v.2.6.0+) with short query parameters (Camacho et al., 2009). Matches covering at least 90% of the spacer and with an e-value < 0.001 were considered to be CRISPR spacer-virus associations. Additionally, virotypes and taxonomically annotated contigs were mapped against the representatives genomes of the viral clusters in the MVP database (Gao et al., 2018) using LAST-959 (Kielbasa et al., 2011). As viral clusters in MVP comprise sequences that have at least 95% identity along at least 80% of their lengths, only matches that fulfill those constraints were kept. The host(s) of a contig was determined from its matching viral cluster.

### Diversity indexes

The Shannon diversity index within-samples (α-diversity) and the Hellinger distance within co-twins (β-diversity) were calculated using *diversity* and *vegdist* functions of Vegan R package for all three abundance matrices generated (function, taxonomy and read recruitment matrices). Correlations between virome α-diversity and microbiome α-diversity were measured using the Pearson correlation coefficient. Correlations between viromes β-diversity and the microbiomes β-diversity was computed with a the Mantel test using the Pearson correlation coefficient. Additionally, the β-diversity between concordant MZ co-twins was compared to the β-diversity between discordant MZ co-twins; p values were calculated using Mann-Whitney U test.

## DATA AND SOFTWARE AVAILABILITY

Jupyter notebooks and scripts describing the data analysis process are available on GitHub at https://github.com/leylabmpi/TwinsUK_virome The sequence data have been deposited in the European Nucleotide Archive under the study accession number PRJEB29491.

## ACKNOWLEDGEMENTS

We thank Laura Avellaneda-Franco for the discussion and contribution in the development of the strategy to remove bacterial contaminants of viromes. This work was supported by grants from the NIH: NIDDK RO1 DK093595 and DP2 OD007444, the Max Planck Society, and the David and Lucile Packard Foundation. The TwinsUK cohort is supported by the Wellcome Trust; European Community’s Seventh Framework Programme (FP7/2007–2013); National Institute for Health Research (NIHR)-funded BioResource, Clinical Research Facility and Biomedical Research Centre based at Guy’s and St Thomas’ NHS Foundation Trust in partnership with King’s College London.

## AUTHOR CONTRIBUTIONS

RL and SPC designed the study. TS and JB were involved in sample collection. SPC and IH generated the data. JLM-G, SPC, JKG, NY, AR and RL analyzed the data. JLM-G, SPC, SCD, AR and RL wrote the manuscript. All authors read and approved the final manuscript.

## DECLARATION OF INTERESTS

The authors declare no competing interests.

## SUPPLEMENTAL INFORMATION LEGENDS

**Figure S1.**
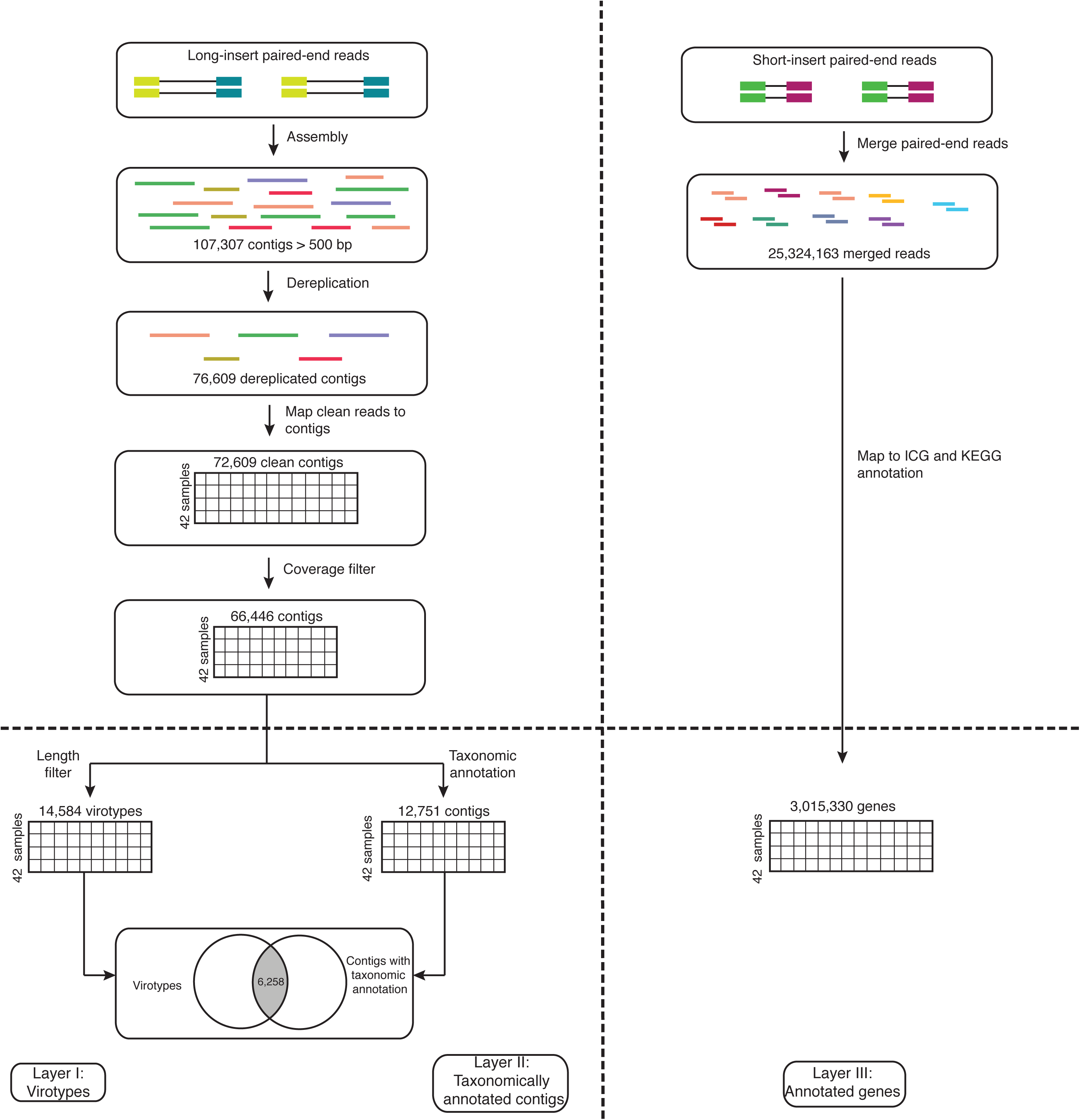
Schematic representation summarizing the procedures applied to **(left)** the “large-insert-size library” and **(right)** the “short-insert-size library” to obtain three different layers of information used to analyze the virome diversity of the microbiome-concordant and microbiome-discordant co-twins.

**Figure S2.**
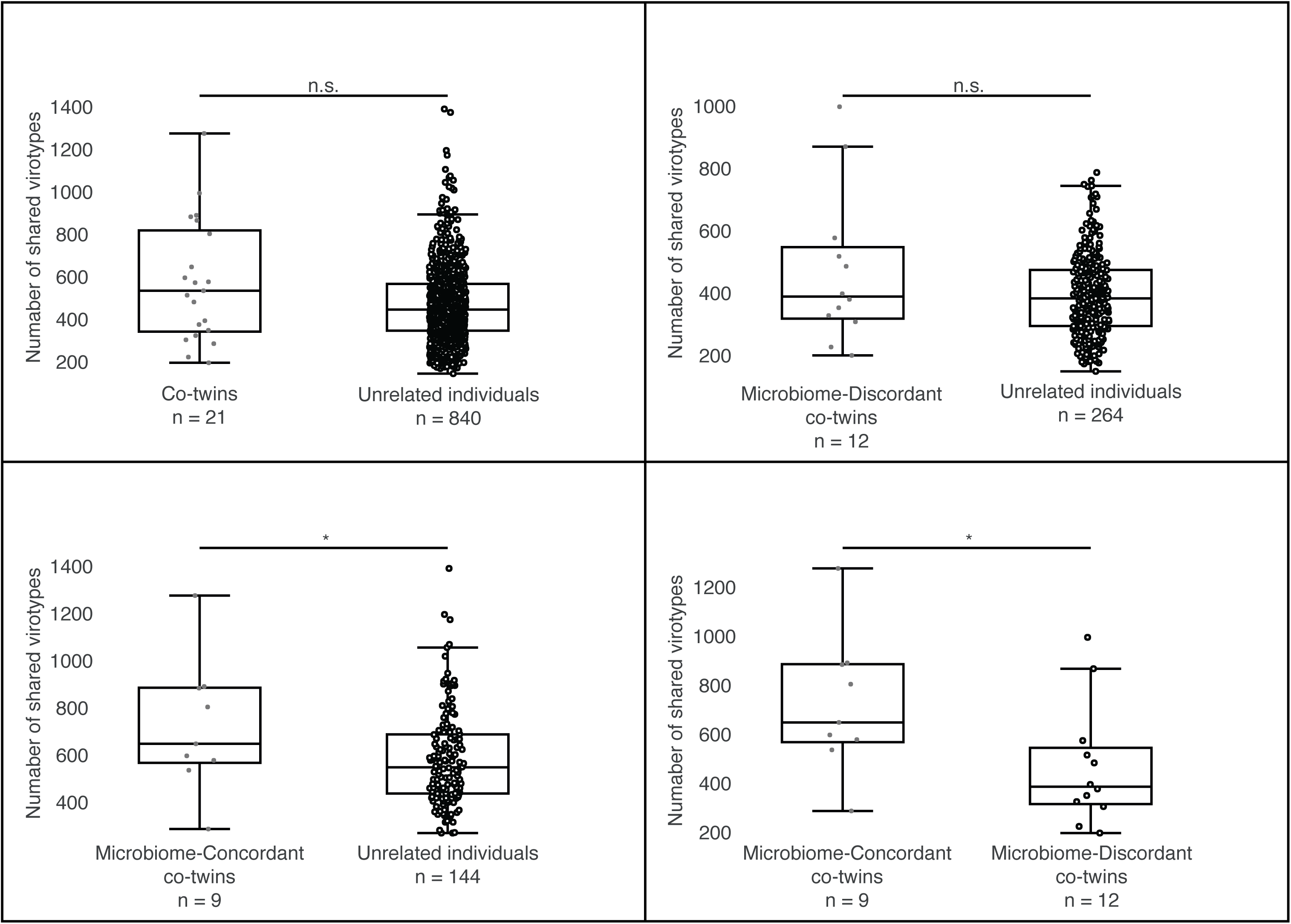
Box plots showing the distribution of the number of shared virotypes between different groups made from the 21 MZ co-twins. (Up left) All co-twins vs unrelated individuals. (Up right) Microbiome-discordant co-twins vs unrelated individuals in the same group. (Down left) Microbiome-concordant co-twins vs unrelated individuals in the same group. (Down right) Microbiome-concordant co-twins vs microbiome-discordant co-twins. Mann-Whitney’s U test. * p < 0.05; n.s: not significant difference.

**Figure S3.**
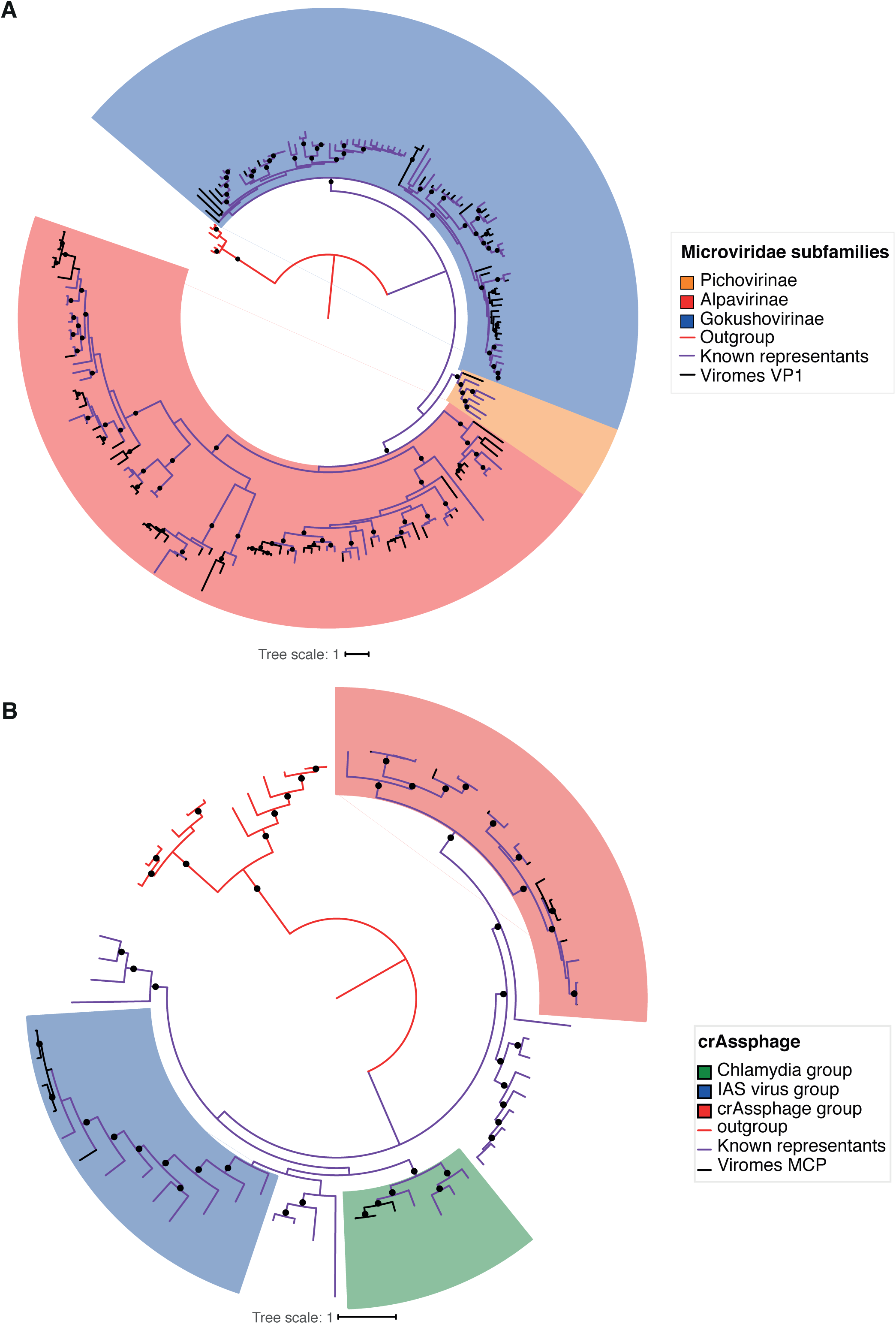
Maximum likelihood phylogenetic analysis of **(A)** the VP1 protein of *Microviridae* phages and **(B)** the MCP protein of crAssphage found in the 42 MZ viromes. Reference sequences are in purple, outgroup sequences are in red while the different MCP or VP1 proteins found in this work are labeled in black. Circles in the nodes indicates bootstrap values above 70%.

**Figure S4.**
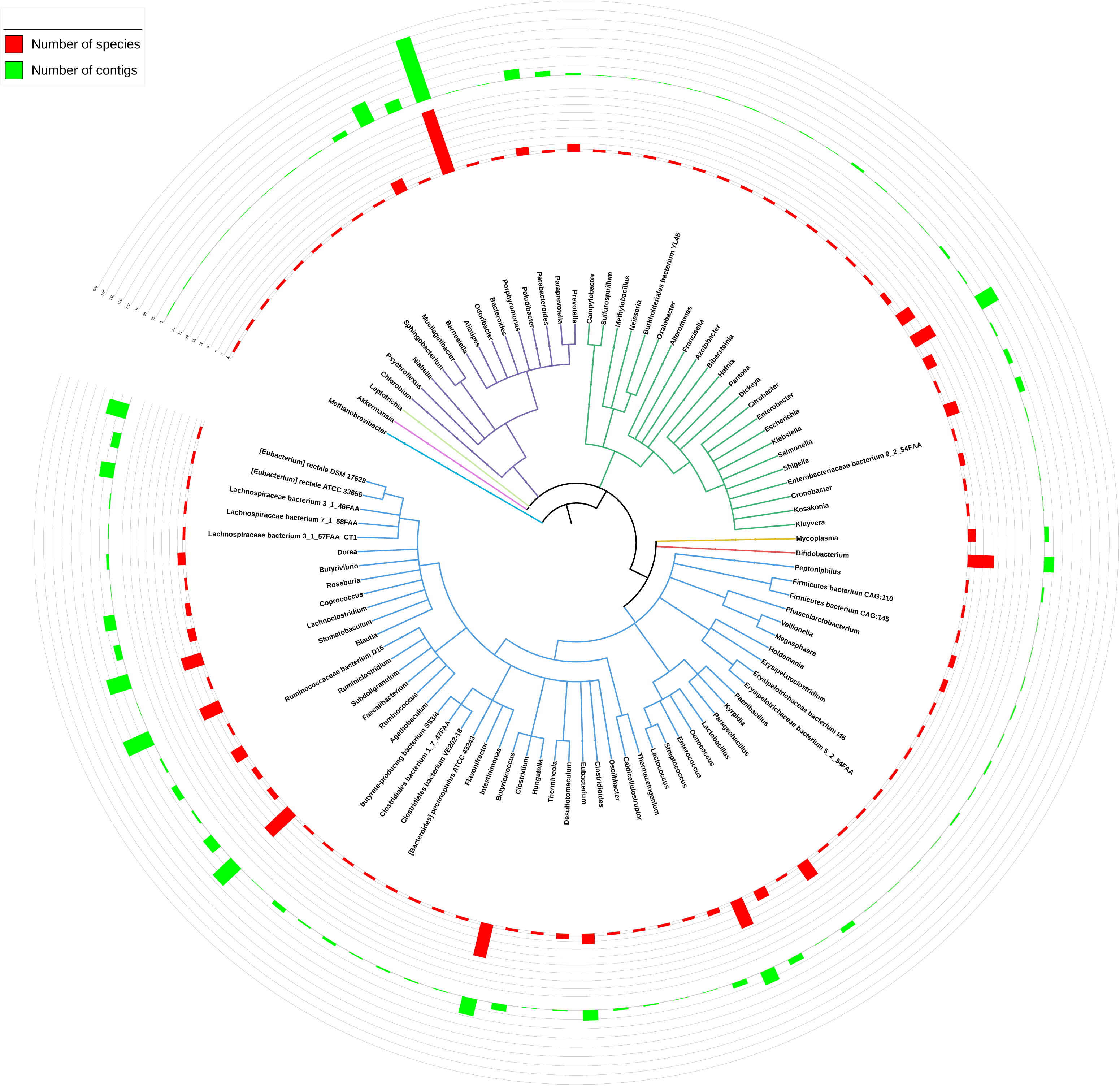
Cladogram based on the NCBI taxonomy showing the bacteria identified as hosts. The cladogram is summarized by genus, and clades are colored by Phylum. Blue: Firmicutes; Red: Actinobacteria; Yellow: Tenericutes; Green: Proteobacteria; Purple: Bacteroidetes; Light green: Fusobacteria; Magenta: Verrucomicrobia; Light blue: Euryarchaeota. Red bars indicate the number of species in each genus, and green bars show the dereplicated number of contigs associated to each genus (i.e. if a contig was found associated to two species in that genus, it is only shown one time).

**Figure S5.**
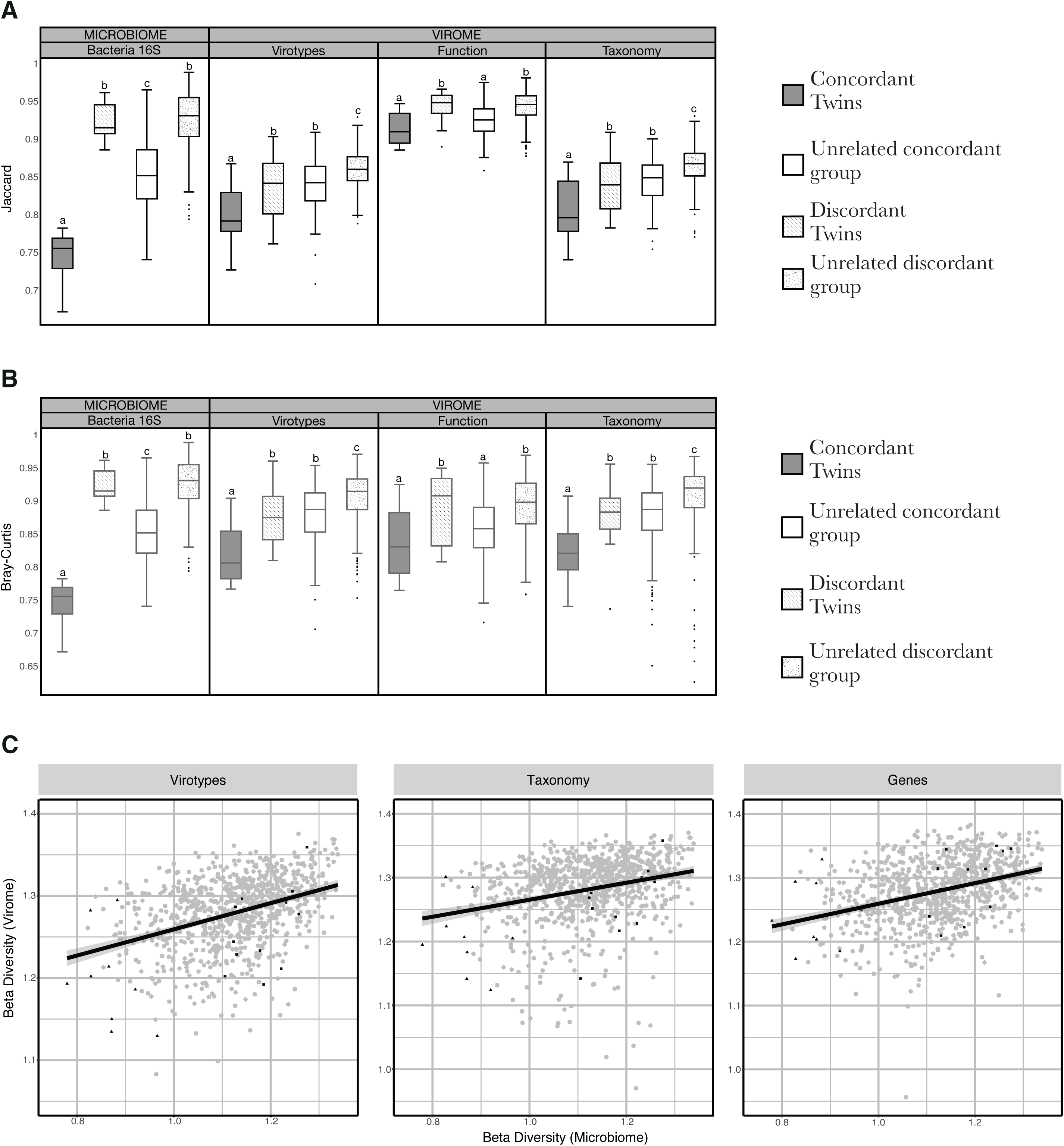
Box plots showing the distribution of **(A)** the Jaccard distances and **(B)** Bray-Curtis distances for microbiomes and viromes, according to the three different layers of information recovered (virotypes, function and taxonomy). Significant differences between means (Mann-Whitney’s U test) are denoted with different letters. Groups and n values as in Figure 6. **(C)** Correlation between virome β-diversity and microbiome β-diversity (n=840). **i) Virotypes:** Pearson correlation coefficient among all individuals = 0.382 (p = 0.0005, Mantel test), m = 0.167, p = 0, R^2^ = 0.157; Pearson correlation coefficient among co-twins = 0.522, m = 0.188, p = 0.015, R^2^ = 0.1508; **ii) Taxonomy annotated contigs:** Pearson correlation coefficient among all individuals = 0.266 (p = 0.003, Mantel test), m = 0.140, p = 0, R^2^ = 0.0796; Pearson correlation coefficient among co-twins = 0.512, m = 0.186, p = 0.017, R^2^ = 0.224; **iii) Genes:** Pearson correlation coefficient among all individuals = 0.344 (p = 0.0009, Mantel test), m = 0.162, p = 0, R^2^ = 0.123; Pearson correlation coefficient among co-twins = 0.53, m = 0.182, p = 0.012, R^2^ = 0.248. Lines describe linear regressions of pairwise distances among all individuals. Triangles indicate concordant-microbiome co-twins and squares indicate discordant-microbiome co-twins.

**Table S1.** Additional information pertaining to the 21 selected MZ twin pairs (metadata), and counts of viromes reads and contigs per sample.

**Table S2.** Median bin coverage of bacterial genomes by VLP reads per sample.

